# Extracting massive ecological data on state and interactions of species using large language models

**DOI:** 10.1101/2025.01.24.634685

**Authors:** François Keck, Henry Broadbent, Florian Altermatt

## Abstract

The contemporary ecological crisis calls for integration and synthesis of ecological data describing the state, change and processes of ecological communities. However, such synthesis depends on the integration of vast amounts of mostly scattered and often hard-to-extract information that is published and dispersed across hundreds of thousands of scientific papers, for example describing species-specific interactions and trophic relationships. Recent advancements in natural language processing (NLP) and in particular the emergence of large language models (LLMs) offer a novel, and potentially revolutionary solution to this persistent challenge, for the first time creating the opportunity to access and extract virtually all data ever published. Here, we demonstrate the transformative potential of LLMs by extracting all types of biological interactions among species directly from a corpus of 83,910 scientific articles. Our approach successfully extracted a network of 144,402 interactions between 36,471 taxa. Performance analysis shows that the model exhibits a high sensitivity (70.0%) and excellent precision (89.5%). Our approach proves that LLMs are capable of carrying out complex extraction tasks on key ecological data on a very large scale, paving the way for a multitude of potential applications in ecology and beyond.

## Main

As stated by Rachel Carson “In nature, nothing exists alone”, it is not the sheer number of species, but rather their interactions and dependencies that define ecological systems. These biological interactions, such as predation, competition, parasitism or mutualism, are the hallmark of ecology and biodiversity sciences. They are a key concept in ecology, and form the foundation of population dynamics, community composition and ecosystem functioning^1–3^. Biological interactions, together with species distribution and functional traits, also represent the key critical information for understanding and responding to increasing environmental challenges^4^ including climate change^5^, biodiversity loss^6,7^, and emerging pathogens^8^.

Yet, data on biological interactions are among the hardest to get hold on. Information on the various types on how species interact is largely based on original natural history observations scattered across a myriad of publications and often dispersed across different research fields. Also, the possible number of interactions is vastly larger than the sum of species involved. Imagine a community of 200 plants and 600 insect herbivores–realistic numbers for even a mid-diversity ecosystem–creating the potential of close to 20,000 pairwise competitive interactions between the plants and close to 60,000 herbivorous interactions alone. Despite only a very sparse part of these potential interactions being realized, assembling and integrating data on species interactions has been a major challenge, with only few food-webs being largely resolved^9,10^, yet alone thinking beyond pairwise interactions^11^.

Because of the central importance of biological interactions for modern ecology and for biodiversity prediction in the current context of global change^11–15^, large database projects have been developed with the aim of collecting machine-readable species-interaction data^16–18^. However, these databases ultimately rely on manual or semi-automatic extraction and integration of data shared by scientists. While data on species occurrences are rapidly increasing and are expected to become less of a limiting factor for ecological research^19^, information on species interactions remains significantly lacking and incomplete^20–22^. This discrepancy arises partly due to the inherent complexity of sampling species interactions^23^ and partly because such data are less systematically recorded and reported.

Integrating and synthesizing ecological knowledge is notoriously difficult. This is largely due to how ecology and natural history findings are disseminated. Typically, they are communicated through scientific articles, which often consist of large volumes of unstructured text^24^. Consequently, the sum of ecological knowledge generated over more than a century, and which we need to mobilize immediately to tackle the global ecological crisis, is dissolved in a vast ocean of text. Unlike structured databases or standardized formats, scientific articles vary widely in their organization. This has made extraction of information laborious and time-consuming or even impossible, such that most information is still hidden (if not lost) in text. Also, ecological research spans diverse subfields and methodologies, resulting in a large and heterogeneous body of literature that can be difficult to navigate and integrate. As a result, ecologists face the daunting task of sifting through a myriad of articles, extracting relevant data, and synthesizing findings into cohesive reviews, databases and meta-analyses. Cataloguing interactions among species started with the very first steps of ecology by Alexander von Humboldt and the tangled bank by Darwin, yet despite generations of ecologists and naturalists collecting and documenting these interactions, they could have been hardly integrated in an inclusive manner.

The current revolution in text mining and natural language processing (NLP) driven by the development of deep neural approaches^25^ and more recently by the advent of large language models (LLMs) is for the first time opening the potential for automated data extraction and synthesis from scientific articles. These models trained on vast amounts of text data, now exhibit unprecedented capabilities in understanding and generating human-like language, allowing to tackle more complex extraction tasks which until now were limited to highly structured and codified textual data (e.g. collections of taxonomic descriptions^26^). In academia the groundbreaking potential of LLMs, including ChatGPT, the popular LLM-powered chatbot by OpenAI, have predominantly been explored and discussed in the context of text generation (e.g. for writing publications^27^) and education (e.g. adoption and use by students and teachers^28–30^), while the capacity of LLMs for information extraction in the form of structured data^31,32^ has received comparatively little attention. First case studies in ecology and biodiversity science use LLMs for automatic data extraction from disease reports^33^, research abstracts^34,35^ and news reports^35^. Nevertheless, these works remain proof-of-concept conducted on small and therefore limited^36^ samples of text, and focusing on specific issues or taxa. Large-scale implementation for more general problems and on a corpus representative of the state of knowledge remains to prove that LLMs can truly revolutionize the synthesis and analysis of ecological knowledge.

Here, we demonstrate the power of LLMs to extract biotic interactions between species directly from the published literature at a very large scale. As shown above, biological interactions are a critically important piece of information for ecological research, and their availability is still very limited. But beyond the hope raised by LLMs to solve this long standing problem, biological interactions are an interesting case study, as they represent a technical challenge for automatic data extraction methods. This is complex information involving two actors (two species) and a directional relationship (the interaction). The algorithm or model has to isolate and qualify these elements, despite the extensive vocabulary and sometimes ambiguous terminology^37,38^ that covers the wide range of biological interactions and the organisms involved. While dictionary-based approaches have been developed to detect biological interactions^39^, their efficiency is strongly limited when confronted with the complexity of the language and the multiplicity of syntactic constructions used in the literature. To date, this complexity precluded any attempt at automated extraction of biological interactions on a large scale.

## Results

From 545,967 processed paragraphs originating from 83,910 scientific articles, the model extracted 649,319 potential biotic interactions. These interactions involved 157,354 unique named entities (theoretically representing names of organisms) among which 70,841 could be taxonomically linked to the NCBI Taxonomy database through TEL. We excluded results from 29 paragraphs for which the model generated an infinite sequence of repeated words and also 4,272 interactions that matched exactly the examples provided in the prompt and were thus most likely the result of an overfit. To merge synonymous nodes and remove potential false positives, we filtered the data to keep only interactions involving taxonomically linked entities. We also merged all edges between two nodes belonging to the same category. This resulted in a final interaction network of 35,471 nodes and 144,402 edges (interactions) originating from 112,647 paragraphs, from 36,044 publications. The interactions were described in the source publications with a very rich and more or less specific vocabulary (14,767 unique labels, see Fig. 2 for an overview of label-to-category associations), demonstrating the importance of a good understanding of the context by the model.

The global set of extracted interactions were filtered and aggregated at different levels of taxonomic resolution (Table 1). For example, the species level network involved 18,589 different species and 46,467 species-to-species interactions distributed in 1,295 components (see Fig. 1 for a detailed view of this network). Interactions were dominated by parasitism (Fig. 1a-c) and the species with the most interactions extracted were species exhibiting a major role in human cultures, such as domestic animals and livestock, or parasites of medical or veterinary interest (Fig. 1d).

**Table 1.** Summary statistics of the extracted network aggregated at different taxonomic levels. The nodes correspond to individual taxa, the edges represent the interactions between the taxa, and the components are the subparts of the main graph that are not connected together.

| **Taxonomic level** | **Nodes** | **Edges** | **Components** |
| --- | --- | --- | --- |
| Mixed (all) | 36,471 | 144,402 | 933 |
| Superkingdom | 4 | 157 | 1 |
| Phylum | 127 | 5,201 | 2 |
| Class | 353 | 9,772 | 2 |
| Order | 1,140 | 26,345 | 11 |
| Family | 4,118 | 42,744 | 58 |
| Genus | 12,980 | 56,404 | 390 |
| Species | 18,589 | 46,467 | 1,295 |

**Figure 1.**
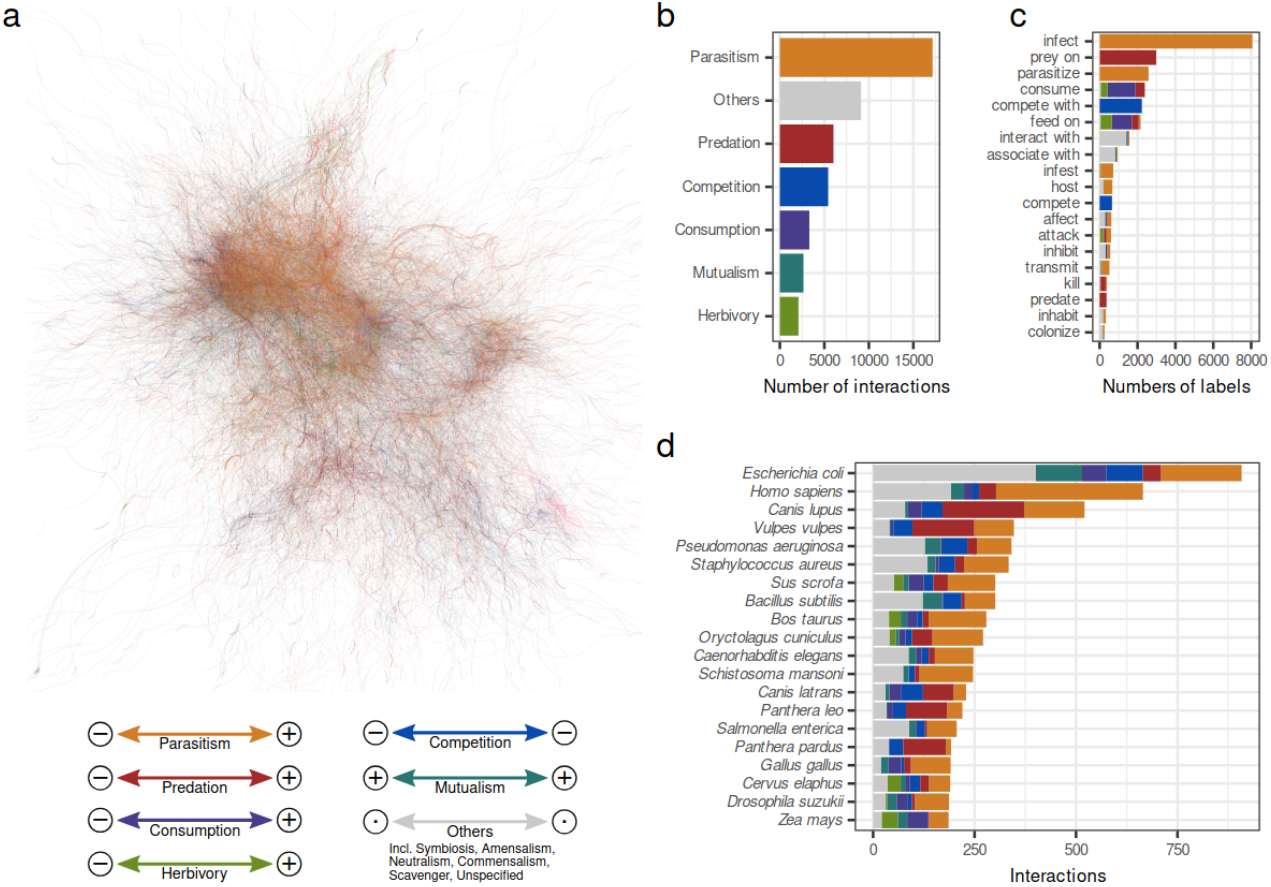
Reconstructed network of interactions at species level. **a**. Overview of the main graph component represented using a force-based (ForceAtlas2) layout. **b**. Number of interactions for the different inferred categories across the whole network. **c**. Number of extracted labels and their inferred categories for the 20 most common labels. **d**. Number of interactions and their inferred categories for the 20 most connected species. In each panel, interaction categories are color-coded as described in the legend, which also indicates for each category how the partners are affected (positively or negatively).

**Figure 2.**
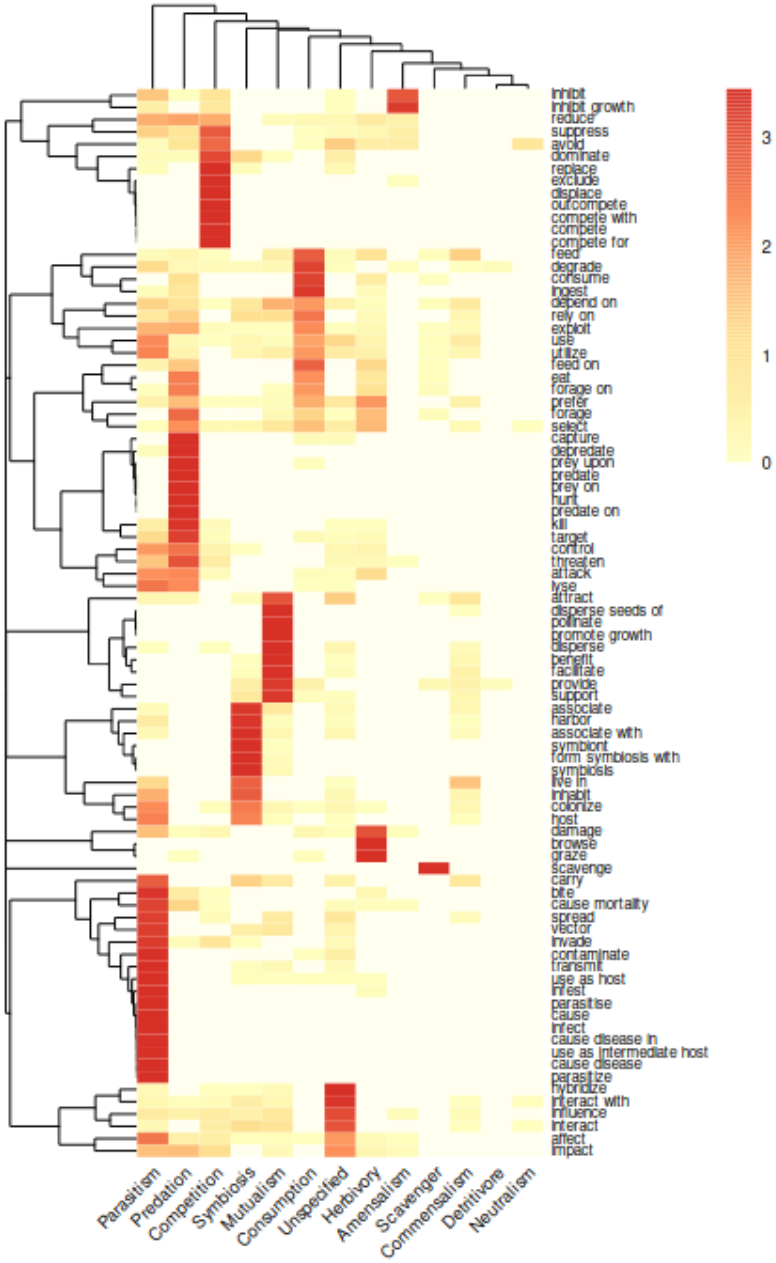
Heatmap showing the association between extracted labels and inferred interaction categories. Values represent the number of times each label has been classified into each category, scaled by row to improve readability. Only the most common labels (extracted more than 250 times) are shown (n = 87). The dendrograms represent the results of hierarchical clustering for rows (labels) and columns (categories).

To compare and validate these LLMs extracted interactions, we manually screened a subset of this text and extracted all interactions based on expert knowledge. The manual annotation of 500 paragraphs in the validation set identified 327 biological interactions. Our automated approach managed to identify 229 of these (true positives), while making 27 errors (false positives). This corresponds to an accuracy of 89.5% and a recall of 70.0%.

## Discussion

As environmental challenges are multiplying and becoming more pressing, it becomes increasingly important to mobilize all available knowledge in ecology. Here, we successfully demonstrated that this is now within reach and made possible by new developments in artificial intelligence and, in particular, the emergence of high-performance large language models. We extracted detailed information on biological interactions, a key parameter in ecology, among thousands of species directly from the scientific literature and at unprecedented scales, with the opportunities to further integrate these data into existing products or to produce original ecological research about the structure and properties of this global ecological meta-network^40,41^. Up to now, access to species interaction data has been one of the most limited and biased, yet one of the most sought after data types^20,22^. These results give us a glimpse of a future where the literature will be scanned on large scales to extract massive amounts of data that were previously beyond the reach of any automated algorithm.

Our study also provides an opportunity to measure the accuracy of automatic LLM extraction in an ambitious and complex case study, and to identify the challenges and limitations of this novel approach when applied at large scale. Our method effectively balanced overall correctness with the ability to identify relevant cases, even in challenging scenarios. While the recall (70.0%) suggests room for improvement in detecting all relevant interactions, the relative high accuracy (89.5%) highlights the robustness of the model in minimizing false positives. Although the extraction relies primarily on a LLM, it is part of a more complex data processing pipeline that guarantees the quality of results. In particular, we implemented a pre-filtering step to target the most relevant documents and a post-filtering step to improve the quality of the results. Pre-filtering is not strictly necessary but allows to restrict the corpus of text, hence limiting the number of requests and therefore reducing the financial and environmental costs of the model. In our experiment, post-filtering proved to be necessary to remove the large number of false positives extracted by the model (e.g. non relevant relationships between an organism and its environment). All these steps, when properly implemented, can be easily automated.

Importantly, as vast and rich as the scientific literature may be, it is inevitably biased, and the data extracted by our approach precisely reflects these biases. The species most represented in our results include humans (*Homo sapiens*), human pathogens (e.g. *Escherischia coli,Pseudomonas aeruginosa,Staphylococcus aureus*) and domestic species (e.g.*Canis lupus,Sus scrofa,Bos taurus*), all of which have been the focus of particular attention from the scientific community. Data extraction can only highlight biases in the literature and does not correct them. This is a factor that must always be considered when examining the results.

A number of challenges and limitations have also been identified regarding the use of proprietary LLMs for scientific research^42^. First, LLMs can pose a problem for reproducibility because outputs can be non-deterministic, i.e., the same query (prompt) might yield different results depending on the model’s internal state. A potential solution is to perform deterministic sampling by using fixed random seeds. This approach makes outputs more consistent across runs but all conditions (hardware, software, model parameters) must be controlled, which is difficult to ensure with cloud-based closed models. Secondly, these models are essentially black boxes. The data and methodology used to train them are generally kept confidential by the developing companies. Such opacity requires researchers to take a step back from the possible biases induced by the methodological choices made by the model’s designers. Analysing the model’s outputs carefully and performing a thorough validation, as done in the present study, is therefore indispensable for any scientific application of this type of approach.

The above challenges are solvable, at least in part, through the use of small open models that are smaller in size but can be optimized and used locally. Despite their smaller size, small-scale LLMs – when appropriately fine-tuned – can achieve impressive results across different applications^43–45^. The advantages of this approach are many^46^: greater independence, greater transparency, better control on reproducibility, and potentially lower costs and environmental imprint. Moreover, the use of a model run locally potentially enables processing documents with more restrictive licenses^47^. However, selection, deployment, optimisation and fine-tuning of such models may require a certain amount of expertise and resources. Despite these challenges, it is clear that the impact of LLMs will be all the greater if they are put directly into the hands of scientists and make their way into laboratories^46,48^. In this respect, small-scale fine-tuned LLM represents a promising direction.

Our approach represents a potentially major paradigm shift with regard to the way we interact with scientific literature, fitting in with the epistemological revolution initiated by the emergence of LLMs^49^. Through our results we show that a new level of qualitative and quantitative synthesis of the available knowledge can be achieved. Building on recently published proof-of-concepts^33–35^, our study grounds the use of LLMs for synthesis sciences and by demonstrating that we can extract accurate structured information from scientific publications on a large scale, it opens the way to a virtually infinite number of applications. Whether for building databases or conducting meta-analyses, the number of questions that can be addressed in ecology is significant. Automated data extraction also paves the way for living syntheses, i.e., studies that are continually updated and whose results and conclusions evolve as new primary results are made available^50^. In the context of the ongoing biodiversity crisis where scientists are confronted with an inflation in the number of publications and the dispersal of information, advancing ecological data access and extraction to a new level will help to better understand and model current and future species distribution and interactions and guide critical management decisions.

## Methods

### Corpus compilation

We screened scientific publications using a large language model to extract all possible biological interactions. The text corpus was sourced from the PubMed Central (PMC) Open Access database, which contains millions of publications spanning various scientific disciplines^51^. To narrow the scope of our analysis, we first refined our corpus to focus specifically on publications related to ecological sciences. Using the OpenAlex database^52^ and its topic classification system, we identified 99,555 publications tagged under the subfield “Ecology” from an initial pool of 6,159,719 PMC publications. We downloaded the corresponding 99,555 XML files from PMC and parsed them to extract 3,552,030 paragraphs for further analysis.

We refined this dataset by retaining only the paragraphs that included at least two distinct taxonomic names and one keyword associated with species interactions. To identify taxonomic names, we employed TaxoNERD^53^, a named-entity recognition (NER) tool utilizing deep neural models for recognizing both scientific and vernacular taxonomic names. TaxoNERD detected a total of 8,160,227 taxonomic names across 1,925,414 paragraphs from 97,681 articles. In parallel, we performed a regular expression-based search to detect species interaction terms using a curated list of 32 relevant keywords. The application of both filters—requiring a minimum of two distinct taxonomic names and one species interaction keyword—resulted in a final corpus consisting of 545,967 paragraphs from 83,910 articles.

## Data extraction

The final dataset of 545,967 paragraphs was subsequently processed using the OpenAI GPT-4o model. Each paragraph was incorporated into a standardized prompt template (Fig. 3), which was then submitted to the OpenAI API for analysis. The prompt template was specifically designed to extract all pairwise species interactions present within the paragraph and structure the output in a tabular format. The output consisted of a four-column table where the first two columns contained the names of the two species involved in the interaction, the third column specified the nature of the interaction as labeled in the text, and the fourth column classified the interaction into its respective category from the list used in the Mangal database^17^. This structured format allowed the systematic analysis and categorization of species interactions within the corpus.

**Figure 3.**
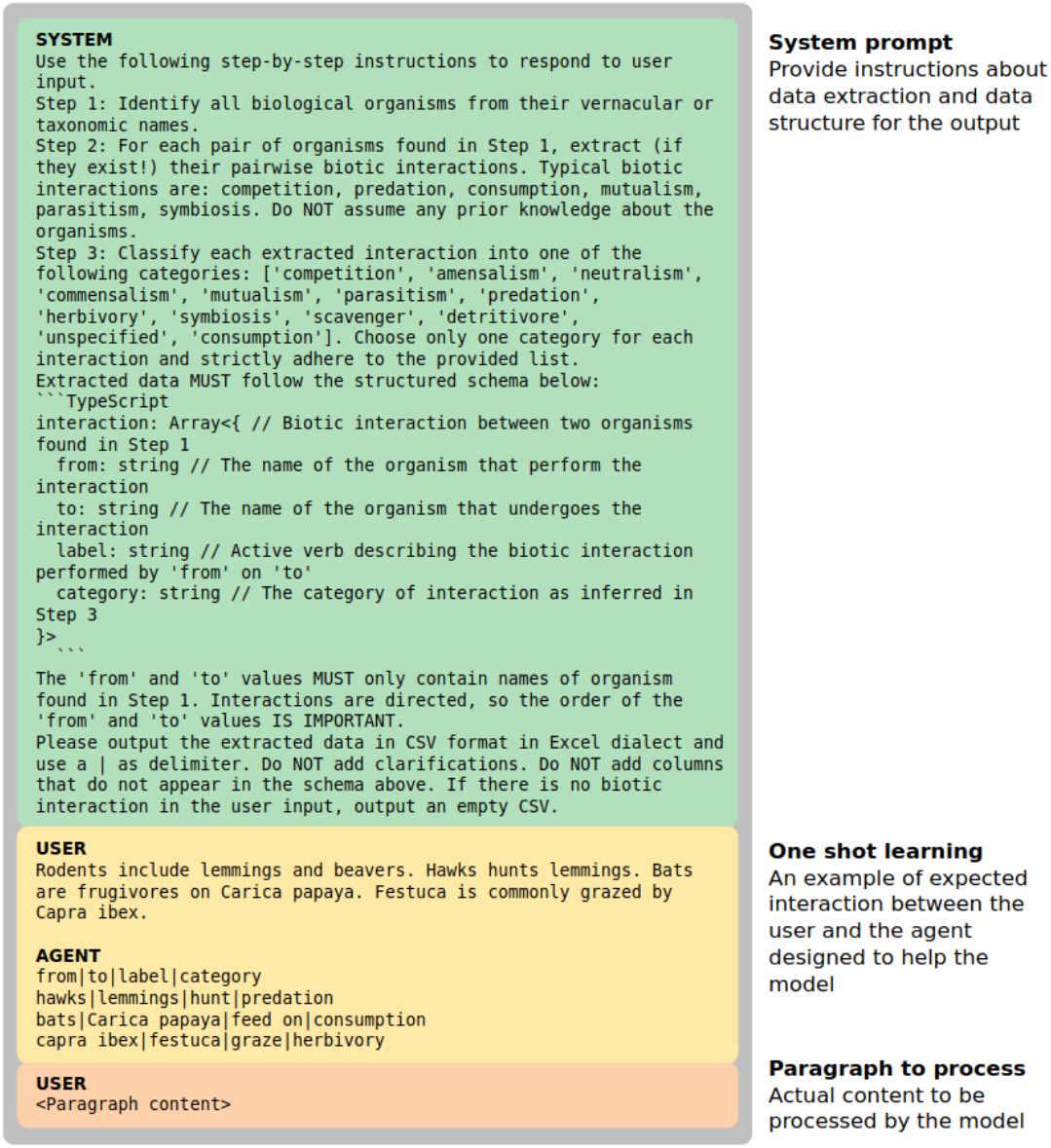
Standardised prompt template used to process each paragraph. First a system prompt is used to give a detailed description of the task to the model. Second, one example of task resolution is provided to the model to refine performances. Finally the content of the actual targeted paragraph is provided for extraction.

To identify interactions involving entities that cannot be linked to living organisms and to harmonize organism names and link them to their upstream taxonomy, we performed taxonomic entity linking (TEL). TEL consists in mapping extracted named entities to corresponding unique identifiers in a target knowledge base, in our case the NCBI Taxonomy^54^. For example, through TEL, the entities dog, domestic dog, *Canis familiaris* and *Canis lupus familiaris* are all linked to the same NCBI Taxonomy ID: 9615 and constitute a unique node in the final network. We implemented TEL through a custom multistep algorithm using data from NCBI, the Encyclopedia of Life (EoL), the Integrated Taxonomic Information System (ITIS) and Wikipedia (see corresponding R script for detailed implementation).

## Performance assessment

To estimate the quality and completeness of the information extracted by our approach, we manually reviewed and annotated 500 paragraphs (≈ 1% of the total corpus) taken at random from all those submitted to the model. Using these human annotated data, we computed two performance statistics. First, the precision, calculated as the fraction of relevant retrieved interactions among all the retrieved interactions and second, the recall, calculated as the fraction of relevant interactions that were retrieved.

## Data availability

The publications used to produce the results are all part of the PMC Open Access Subset collection (https://www.ncbi.nlm.nih.gov/pmc/tools/openftlist/). All data extracted are openly available at: https://github.com/fkeck/gpt_interactions

## Code availability

All code used for analysis is available on GitHub: https://github.com/fkeck/gpt_interactions

## Acknowledgments

Funding is by the University of Zurich “Fonds zur Förderung des Akademischen Nachwuchs” (to FK) and the Swiss National Science Foundation (grant 310030_197410) to FA.

